# Interaction with complement factor C1q can contribute to and improve monoclonal antibody-mediated protection against lethal HSV-1 challenge

**DOI:** 10.64898/2026.10.19.760222

**Authors:** Lesle M. Jimenez, Matthew D. Slein, David A. Leib, Margaret E. Ackerman

## Abstract

Herpes simplex virus (HSV) has extensive strategies to evade host defense, including mechanisms that limit clearance by the complement cascade. Using Fc engineering, we incorporated a complement-enhancing (E430G, EG) or a complement-abrogating mutation (K322A, KA) to a panel of glycoprotein D- and glycoprotein B-specific monoclonal antibodies (mAbs) to assess the role of complement-driven antiviral activity in a neonatal mouse HSV-1 infection model. *In vitro* characterization confirmed that Fc modifications preserved antigen recognition while selectively amplifying or reducing complement activity. *In vivo*, complement modulation altered the protective capacity of select mAbs to protect against lethal HSV-1 challenge, demonstrating that complement contributes to antibody-mediated control of HSV-1 infection in an antibody- and antigen-dependent manner. The differential effects observed across the mAb panel indicate that complement activity does not function as a uniform determinant of antibody efficacy, but instead operates in concert with antibody-specific properties to shape antiviral protection. Collectively, these findings establish complement-mediated activity as an important component of antibody function in HSV-1 infection, and highlight Fc engineering as a strategy to optimize antibody-mediated protection. More broadly, these findings underscore the multifaceted nature of antibody-mediated protection and provide a mechanistic framework for considering complement-dependent functions in the development of antibody-based therapeutics.

## Introduction

Viruses have evolved successful strategies to evade host immune defenses. As the immune system deploys both innate and adaptive mechanisms to recognize and clear viral infections, infectious pathogens in turn adapt and develop countermeasures to evade host defenses. As a major component of innate immunity, the complement system acts as a first line of defense that readily responds to viral infections. It is composed of more than 50 proteins, both soluble and membrane-bound, that engage in a sequential cascade of reactions culminating in pathogen clearance^1,2^. The alternative, classical, and lectin pathways, which are activated in different contexts, converge upon the C3 convertase, ending in the common terminal pathway^3,4^. Activation of the complement system results in: phagocytosis of pathogens, promotion of inflammation, and lysis of pathogens or infected cells via membrane attack complex (MAC) formation^5^. Thus, this system is crucial for immune defense and surveillance.

The broad and immediate immunity induced by the complement system is conserved throughout evolution. This ancient defense mechanism predates the adaptive immune system and has been found to be present across a variety of species^6,7^. Phylogenetic conservation can be found in invertebrates, including Cnidaria, and components are present in all of deuterostomes (fish, insects, reptilians, mammals)^1^. Relevant to the complement activity of antibodies, C1q predates the development of the humoral immune system, which evolved later to leverage the cascade through pathogen-specific immunoglobulins G and M. In sum, whether reinforced through connections to adaptive immunity or not, the complement system is a vital component of host defense.

In the context of viral infections, complement exhibits various antiviral activities. Following activation, all three pathways lead to complement deposition on the virion, which can result in blocked interaction with host cell receptors and aggregation of virus particles^8^. The terminal components of complement form MAC on the membrane of infected host cells and virions, resulting in lysis^9^. Opsonins resulting from the complement cascade can bind viral membranes and infected host cells, tagging them for engulfment via phagocytic cells with complement receptors^10^. Complement activation also produces anaphylatoxins are potent chemoattractants for effector cells, recruiting neutrophils and monocytes to the site of infection and amplifying the inflammatory response^5,11^. Though commonly thought of as a non-specific first line of defense, the complement system is unique in its ability to bridge innate and adaptive immune responses. Complement receptors CR1 and CR2 are found on B cells and follicular dendritic cells, where they can contribute to antigen presentation and reduce the threshold for B cell activation and stimulate antigen-specific antibody responses^9,12^. Complement also helps promote formation of germinal centers and prolong B cell expansion. Thus, complement is uniquely positioned to provide robust innate responses while enhancing adaptive responses as well.

Given its conservation across species and the critical role it plays as a first line of defense, complement is a target of viruses. The selective pressure exerted by the complement system has driven viruses to evolve to evade or dysregulate complement. These include: encoding proteins that mimic complement proteins/regulators, inhibiting complement activation, using complement receptors for cellular entry, and incorporating host complement regulatory proteins for viral particle defense^13^,^10^. Herpesviruses have a variety of evasion molecules that limit the antiviral activity of antibodies and complement. In particular, herpes simplex virus (HSV) actively inhibits complement activation via glycoprotein C (gC) expressed on the viral membrane. gC binds complement protein C3b and C3b fragments^14–16^, thus inhibiting downstream complement activity and altering complement-mediated antiviral activity. This inhibitory activity is especially important during infection when the host immune system generates antibodies against HSV. These antibodies, once bound to their target, can activate the classical cascade via C1q binding to IgG or IgM Fc domains. The complement cascade will proceed until it reaches the C3 convertase that generates C3b molecules. Once gC binds C3b, it essentially inactivates the cascade and prevents formation of the C5 convertase, regardless of which complement pathway was activated. Two important sites on gC allow it to interfere with complement: a C3b binding domain (found in both HSV-1 and 2) and a C5/properdin-interacting domain (HSV-1 only)^16,17^. Compared to wildtype virus, gC-mutant viruses are less virulent *in vivo*^17–19^ and readily neutralized *in vitro*^20–22^. Thus, gC plays a critical role in immune evasion and protects HSV-1 from complement-mediated neutralization and lysis of virions and infected cells.

To this end, understanding how antibody-complement interactions affect antiviral activity and whether modulating such interactions can enhance antibody efficacy may prove valuable to the development of an effective prevention and treatment strategies. In this work, we utilized a panel of monoclonal antibodies (mAbs) specific for glycoproteins D (gD) and B (gB), which serve as immunodominant attachment and fusion proteins, respectively. These mAbs provide protection in preclinical models^23–31^, and effector functions, including those mediated through Fcγ receptors (FcγR), are known to contribute in some cases^23,24,32^, but the mechanism(s) responsible for mediating protection remain to be fully defined. Extending the mechanistic profiling of this well-characterized panel of mAbs promises to elucidate the balance of contributions from different effector functions and inform on how distinct components of antibody-mediated host defense might collaborate in combating HSV-1, a highly evolved and evasive virus.

## Results

### Incorporating Fc modifications to manipulate antibody-complement interactions

To determine how complement contributes to antibody-mediated protection against HSV-1, we constructed a panel of HSV-specific mAbs with mutations that either enhance or abrogate complement activity (**Figure 1A**). The E430G (EG) mutation in the CH3 Fc domain enhances antibody-complement interactions by increasing the hexamerization propensity of IgG, thus favoring binding of C1q and subsequent complement activation^33^. This oligomerization is intended not to occur spontaneously, but rather to be antigen-dependent, requiring mAb to be present at a high local concentration, such as occurs at the surface of a virion or virus-infected cell. In contrast, the K322A (KA) mutation abrogates complement activity by disrupting ionic interactions in the CH2 Fc region where C1q binds, thus reducing complement activation by disfavoring initial C1q binding^34^. Both complement variants have been reported to retain FcyR binding^33,34^. With these engineered antibodies containing complement-related mutations, subsequent comparisons with unmodified human IgG1 (WT) can be made to determine the role of complement in antibody-mediated antiviral activity.

**Figure 1:**
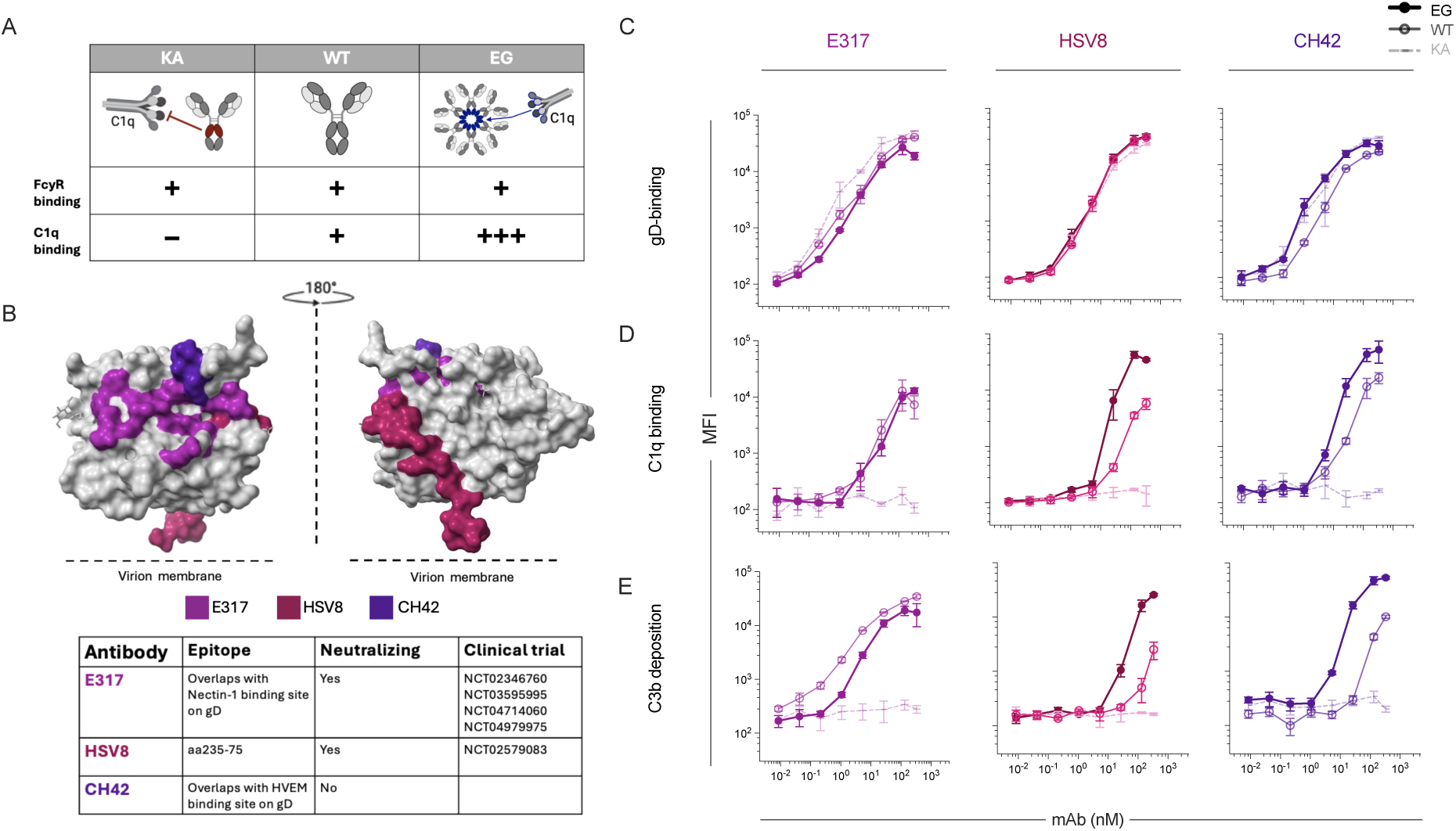
Fc-engineering modifications to modulate antibody interactions with complement C1q. **A.** Schematic and table depicting KA and EG Fc mutations and effects on interactions with FcγR and C1q. **B.** Crystal structure of glycoprotein D (PDB: 1JMA) with mAb epitopes (E317 PBD: 3W9E; CH42 from [Wang el al]; HSV8 from [De Logu et al]) indicated in color. Table presents epitope, neutralization activity, and clinical trial identifiers. **C-E.** Functional characterization of gD mAbs E317 (left), HSV8 (center), and CH42 (right) in EG, WT, and KA forms in dark thick, medium thin, light dotted lines, respectively. Antibodies were titrated and assessed for antigen binding (**C**), C1q binding (**D**), and C3b deposition (**E**). Graphs depict median fluorescent intensity (MFI). Error bars indicate standard error of the mean across replicates.

Three human mAbs specific to gD, the viral protein responsible for attachment to host receptors Nectin-1 and HVEM^35^, were studied in this work (**Figure 1B**). Both E317^36^, which blocks both the HVEM and nectin-1 receptor binding sites, and HSV8^37^, which has yet to be structurally resolved, potently neutralize HSV *in vitro*, whereas CH42 is a non-neutralizing mAb specific to the HVEM binding site^28^. All can provide at least partial protection against lethal HSV infection in both adult and neonatal mouse models, though they differ in efficacy in association with neutralization potency and other factors^25,28,32,37^. The role of complement in protection mediated by these mAbs is not known.

The EG and KA variants of each mAb bound to recombinant gD extracellular domain equally well as the corresponding WT form (**Figure 1C**). In contrast, the KA variants of each mAb exhibited no ability to subsequently be recognized by purified C1q (**Figure 1D**), or to drive deposition of C3b in the presence of complement active serum (**Figure 1E**). The EG variants of HSV8 and CH42 exhibited enhanced C1q binding (**Figure 1D**) and C3b deposition (**Figure 1E**), but the EG variant of E317 did not show reliably potentiated C1q or C3b ligation. Thus, initial characterization of modified gD-specific mAbs confirmed preserved antigen binding, effective elimination of complement cascade initiation by the KA variants, and the expected enhancement afforded by the EG mutation for HSV8 and CH42.

However, the degree of C1q and C3b ligation afforded by WT and EG mAbs in these *in vitro* assays was dependent on antigen density. Under higher antigen density conditions, while the KA variant of each mAb still neither bound C1q nor drove C3b deposition, the relative activity of WT and EG mAbs differed (**Supplemental Figure 1A**). In this case, both E317 and HSV8 EG and WT forms exhibited similar complement cascade initiation, while a substantial increase in both C1q binding and C3b deposition was afforded by CH42 EG as compared to WT. The Fc engineered mAbs generally exhibited the expected profile; specific assay conditions, however, substantially impacted their apparent relative activity, as in other reports characterizing complement activation *in vitro*^38–41^.

To extend our investigation to include the viral fusion protein, four gB-specific antibodies were also expressed in unmodified, EG, and KA forms (**Figure 2A**). Human antibodies Hu2c and HDIT102^31^ bind to overlapping epitopes in domain I that are relatively conserved in position between pre- and post-fusion conformations of gB^31,42^. Antibody D48^42–44^ prefers the post-fusion conformation, but can bind gB in both states. In contrast, the epitope of the non-neutralizing humanized (murine chimeric) antibody BMPC-23^45^ includes amino acid residues that are exposed in the post-fusion conformation but buried in the pre-fusion state. Again, each of these mAbs exhibit varying *in vitro* neutralization potencies, and *in vivo* protection in adult and neonatal HSV challenge models^24,45^.

**Figure 2:**
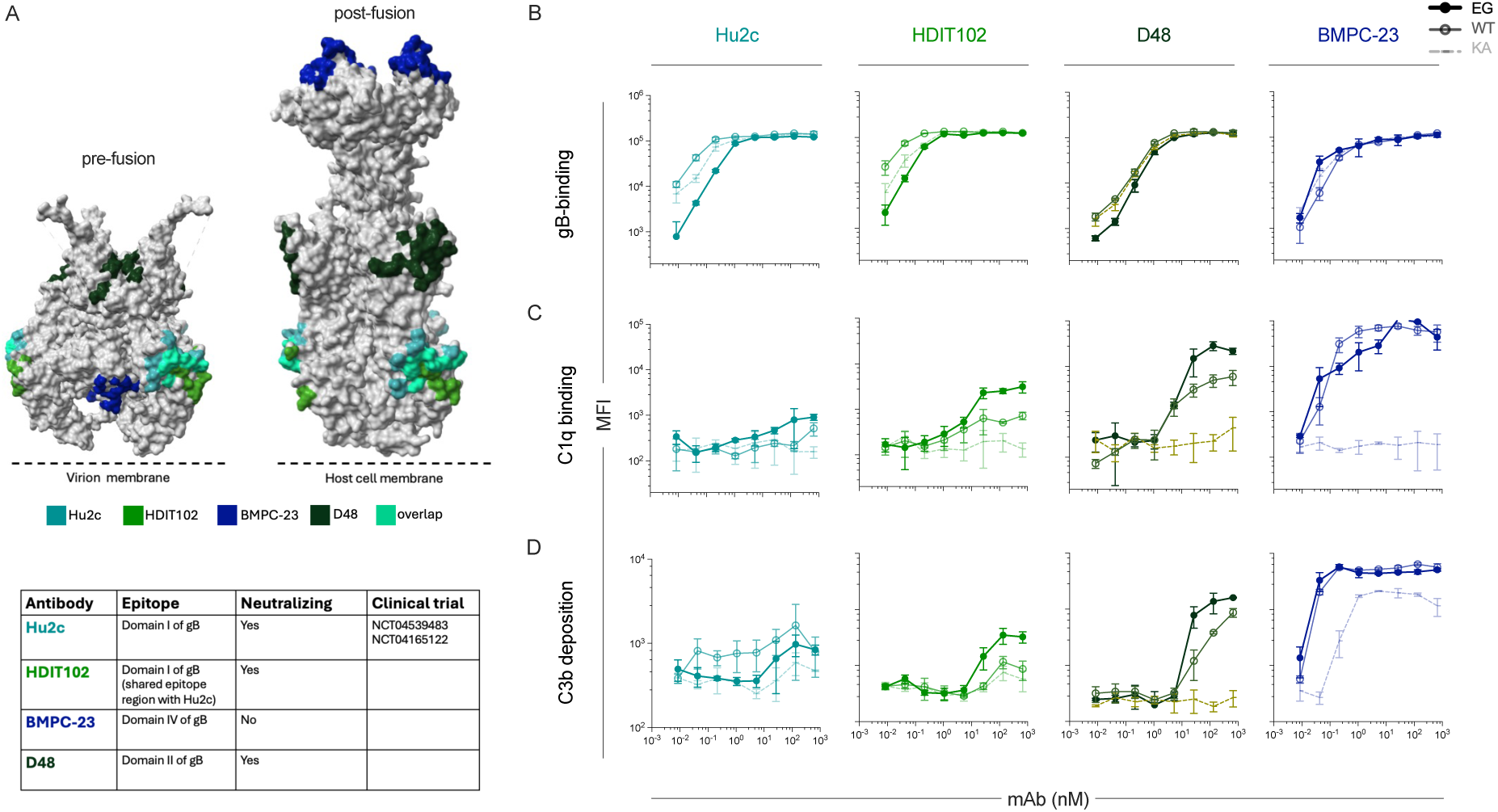
Characterization of gB-specific mAbs. **A.** Crystal structures of the pre- (left, PDB: 6z9m) and post- (right, PDB: 2gum) fusion structures of the gB trimer with epitopes for each mAb indicated in color. Table presents epitope, neutralization activity, and clinical trial identifiers. **B-D.** Functional characterization of gB mAbs Hu2c (left), HDIT102 (center left), D48 (center right), and BMPC-23 (right) in EG, WT, and KA forms in dark thick, medium thin, light dotted lines, respectively. Antibodies were titrated and assessed for antigen binding (**B**), C1q binding (**C**), and C3b deposition (**D**). Graphs depict median fluorescent intensity (MFI). Error bars indicate standard error of the mean across replicates.

For each gB-specific mAb, WT IgG1 and Fc variants bound similarly to recombinant gB protein, suggesting that Fc domain modification did not impact antigen recognition (**Figure 2B**). Despite similar binding to antigen, the KA variant of each antibody generally failed to show either C1q binding (**Figure 2C)**, or C3b deposition (**Figure 2D**). The sole exception to this rule was presented by the KA form of BMPC-23. Despite undetectable C1q binding, this humanized antibody with murine variable domains exhibited a reduced but clearly detectable ability to drive C3b deposition in this assay relative to its WT and EG forms.

In contrast, a detectable increase in C1q binding was seen for the EG variants of Hu2c, HDIT102, and D48 (**Figure 2C**). Again, BMPC-23 proved to be an exception, as EG and WT forms of BMPC-23 exhibited similar C1q binding levels. Consistent with C1q binding results, increases in C3b deposition were seen for the EG variants of HDIT102 and D48 but not Hu2c or BMPC-23 (**Figure 2D**). Under conditions of higher antigen density, again, the KA variants of each tested mAb remained inactive, and WT and EG forms of Hu2C and HDIT102 exhibited similar activity, while the activity of BMPC-23 was better potentiated in EG relative to WT form (**Supplemental Figure 1B**).

Overall, while the profiles of individual mAbs differed in terms of antigen binding and C1q and C3b deposition activity, potentially in association with their distinct characteristics, their epitope-specificities, or to factors associated with assay design, Fc engineering of gD- and gB-specific mAbs generally resulted in the intended complement modulation while retaining comparable antigen binding. Consistent with other studies characterizing the complement-activating properties of antibodies *in vitro* ^46–50^, the extent of functional enhancement varied across mAbs and testing conditions. Here, the EG modification more substantially improved the complement activity of both non-neutralizing mAbs than their neutralizing alternatives, particularly in the presence of high-density antigen.

### Impact of complement interactions on mAb neutralization potency in vitro

Because complement has been reported to enhance antibody-driven viral neutralization of HSV as well as other viruses^51–55^, we next assessed mAb neutralization potency in the presence and absence of complement. In the absence of complement, gD-specific mAbs, E317 and HSV8 potently neutralized HSV-1 strain 17^56^ in a plaque reduction assay while CH42 did not (**Figure 3A**). Consistent with the results of antigen binding experiments, EG and KA variants of each mAb exhibited neutralization profiles similar to their unmodified WT counterparts, demonstrating that direct antiviral activity was not impacted by Fc modification.

**Figure 3.**
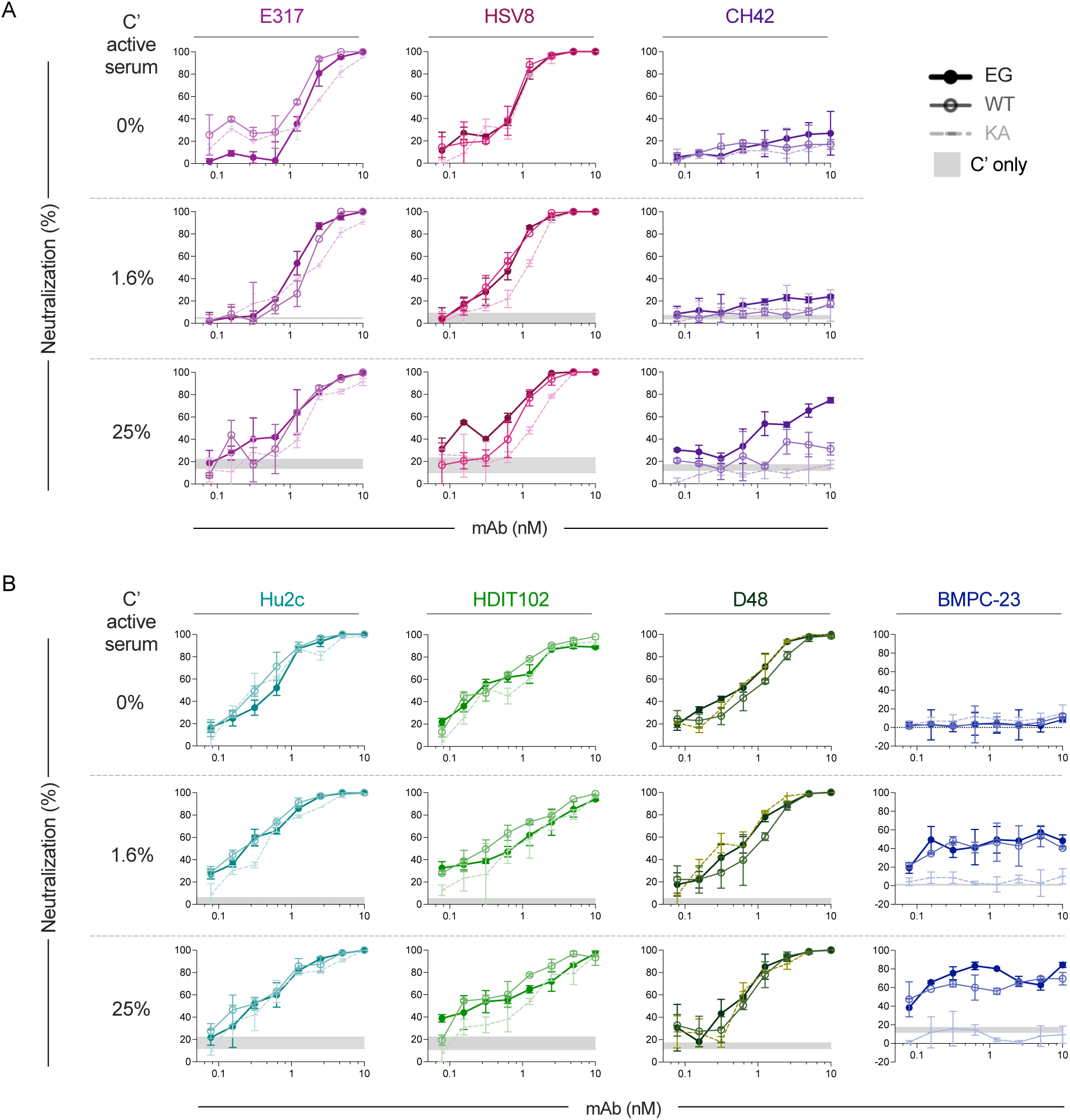
Complement does not add a significant benefit to neutralizing activity of all gD- or gB-specific mAbs. **A.** Neutralization potency of gD mAbs E317 (left), HSV8 (center), and CH42 (right) in EG, WT, and KA forms in dark thick, medium thin, light dotted lines, respectively. Antibodies were titrated and assessed for neutralization in media alone (top) or the presence of complement active serum at a low (center) and high (bottom) concentration. **B.** Neutralization potency of gB mAbs Hu2c (left), HDIT102 (center left), D48 (center right) and BMPC-23 (right) in EG, WT, and KA forms in dark thick, medium thin, light dotted lines, respectively. Antibodies were titrated and assessed for neutralization in the presence of complement at a high (top) and low (middle) dilution, and in the absence of complement (bottom). Active complement serum (without mAb) was included as a negative control for baseline complement-mediated activity. Error bars indicate standard deviation.

Complement-dependent enhancement of neutralization was assessed by conducting the plaque reduction assay with the addition of guinea pig serum as a source of active complement, which enhanced the neutralization activity of pooled polyclonal human serum from seropositive donors (**Supplemental Figure 2**). At either low (1.6%) or high (25%) levels of complement-active serum, KA, WT, and EG variants of E317 and HSV8 neutralized relatively similarly to each other and to these mAbs in the absence of complement (**Figure 3A**). In contrast, while dilute complement-active sera had no impact on CH42, higher concentrations elevated the direct antiviral activity of this typically non-neutralizing mAb to above the level observed following addition of complement-active sera alone (**Figure 3A**). This increase was greatest for the EG variant, intermediate for WT, and not observed for the KA variant.

Similar results were observed for gB-specific mAbs. In the absence of active complement, Hu2c, HDIT102, and D48 potently neutralized HSV-1 while BMPC-23 did not (**Figure 3B**). Again, consistent with their equivalent antigen binding, KA, WT, and EG variants neutralized similarly. Neither dilute nor more concentrated complement-active sera influenced Hu2c, HDIT102, or D48 neutralization potency. WT and EG forms of BMPC-23, by contrast, which failed to neutralize in the absence of complement, reduced plaque numbers modestly in its presence (**Figure 3B**). While EG was not enhanced relative to WT, the specificity of this observation was supported by the lack of enhancement observed for the KA variant.

Overall, only the non-neutralizing mAbs exhibited an increase in direct antiviral activity in the presence of complement. This enhancement was specific for complement-binding mAbs (EG and WT) and was not observed for the C1q-binding knockout KA. It also presented for both gB- and gD-targeting non-neutralizing mAbs, demonstrating that the ability to ligate C1q is necessary for enhancement of direct antiviral activity in the presence of complement-active sera. In contrast, complement-active sera did not alter the neutralization potency of neutralizing antibodies in this panel, regardless of their antigen target or epitope. Collectively, this data provides for mechanistic dissection of the impact of C1q binding capacity on mAb activity.

### The role of complement activation in mAb-mediated protection from HSV-1 challenge

To define the role of antibody-mediated mechanisms of complement-driven antiviral activity *in vivo*, 2-day old C57BL/6J pups were administered mAb intraperitoneally and immediately challenged intranasally with a lethal dose 1x10^4^ plaque forming units (PFU) of HSV-1 (**Figure 4**). Antibody dose was selected with the goal of observing roughly 50% survival for each mAb in WT form, which would, in theory, provide similar power to evaluate both whether eliminating C1q binding reduced protection, as whether enhancement of this activity increased protective efficacy. At this dose, all WT forms of gD-specific mAbs, whether neutralizing or not, provided survival benefit as compared to the HIV-specific mAb VRC01^57^ evaluated as an isotype control (WT IgG1) (**Figure 4A-C**).

**Figure 4:**
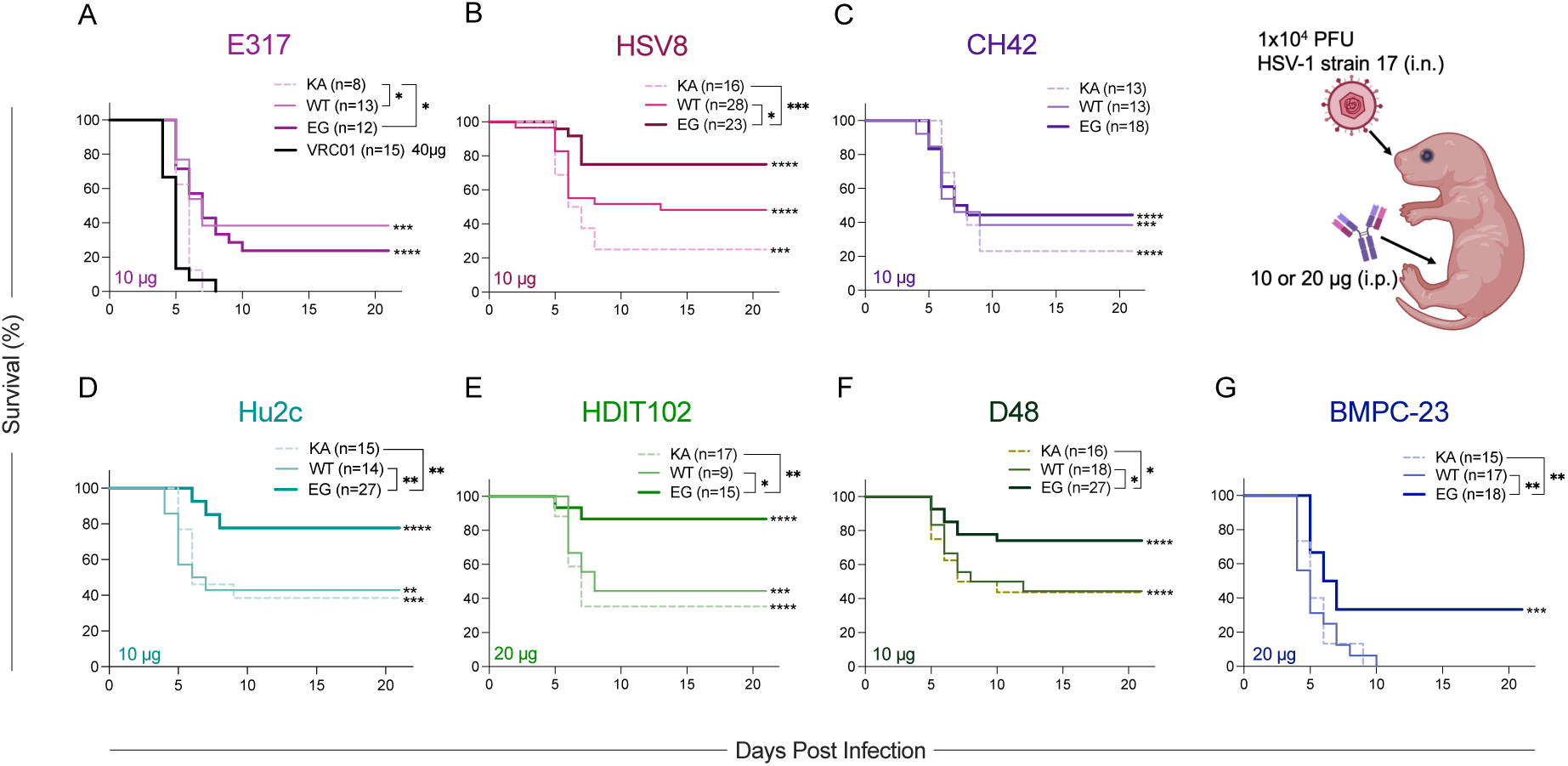
Complement activity contributes to and can improve protection from lethal HSV-1 challenge. **A-F**. Protection afforded by mAb treatment. HSV-specific mAbs were delivered intraperitoneally (i.p) to 2-day old C56BL/6J pups immediately before lethal intranasal (i.n.) challenge with 1x10^4^ plaque forming units (PFU) of lab-adapted HSV-1 strain 17. Survival of pups given gD-specific mAbs E317 (**A**), HSV8 (**B**), or CH42 (**C**), and gB-specific mAbs: Hu2c (**D**), HDIT102 (**E**), D48 (**F**), or BMPC-23 (**G**) at indicated dose. WT, KA, or EG variants are shown in in dark thick, medium thin, light dotted lines, respectively. Statistical significance compared to WT VRC01 (40 μg) isotype control (black, panel **A**) as determined by the log-rank (Mantel-Cox) test is shown on line graphs, and inset reports the number of animals per group and statistical significance between WT, KA, and EG mAbs (*p < 0.05, **p < 0.01, ***p < 0.001, ****p < 0.0001).

In contrast, and despite its high neutralization potency, pups treated with the KA variant of E317 were no more likely to survive lethal challenge than those receiving isotype control (**Figure 4A**). Additionally, while protection was afforded by both WT and EG, treatment with either of these variants led to similar rates of survival, consistent with the lack of enhanced function of E317 EG *in vitro*. Patterns differed for HSV8, the other neutralizing gD-specific mAb. Pups treated with the EG variant exhibited improved survival compared to WT. While ∼20% fewer pups survived when treated with KA than WT HSV8 (p = 0.14), this decrease failed to meet the nominal threshold for statistical significance, and protection was still greater than the isotype control (**Figure 4B**). Lastly, for CH42, KA, WT, and EG provided similar levels of protection (**Figure 4C**), suggesting that complement activation capacity played no role in protection afforded by this non-neutralizing gD-specific mAb.

The gB-specific mAbs presented more uniform results. At the concentrations evaluated, WT forms of Hu2c (**Figure 4D**), HDIT102 (**Figure 4E**), and D48 (**Figure 4F**) but not BMPC-23 (**Figure 4G**) provided protection from lethal challenge. For each of these mAbs, the KA variant provided equivalent protection (**Figure 4D-G**), suggesting that ablating complement activity does not reduce the efficacy of gB-specific mAbs. Despite this apparent independence of complement activation, for all gB-specific mabs in this panel, pups treated with the complement-enhancing EG variant had significantly increased probability of survival compared to their WT-treated counterparts (**Figure 4D-G**). This enhancement was observed for both neutralizing and the non-neutralizing mAb, and regardless of epitope, demonstrating a generalized benefit to result from enhancement of antibody-complement interactions.

Overall, loss of C1q binding reduced the efficacy of two of three mAbs, and enhancing C1q ligation improved the efficacy of one of the three gD-specific mAbs. In contrast, the efficacy of all four gB-specific mAbs was improved by augmenting but not reduced by diminishing complement interactions. That complement abrogating and enhancing mutations exhibit context-dependent impacts *in vivo* may not be surprising given their sensitivity to context *in vitro*; these activities may vary due to antibody characteristics such as antigen specificity and affinity, binding epitope and valency, and neutralization potency, among other factors. Thus, while the impact of Fc modification differed across the panel of mAbs, these results collectively support an important role for complement in antibody-mediated antiviral activity *in vivo*.

### Clinical isolates of HSV-1 are susceptible to antibody-dependent complement activity

The prior experiments were conducted with a lab-adapted virus strain; clinical isolates may present different resistance or sensitivity to complement cascade activation induced by mAbs, select mAbs were tested for efficacy against lethal challenge with the clinical isolate HSV-1 NS^58^. As previously described, 2-day old C57BL/6J old pups were injected intraperitoneally with mAb and immediately challenged intranasally with 1x10^4^ PFU of HSV-1 NS. Here again, the dose of each mAb was selected in order to support detection of both increases and decreases in efficacy relative to WT.

All three variants of the three mAbs selected on the basis of exhibiting sensitivity to complement activity-altering mutations (HSV8, Hu2c, HDIT102) provided protection against HSV-1 compared to a higher dose of the VRC01 isotype control (**Figure 5**). For the gD-specific mAb HSV8, pups treated with the EG variant had increased survival compared to those treated with IgG1, while the KA variant protected similarly to WT (**Figure 5A**). For the gB-specific mAb Hu2c, WT and the EG variant provided equivalent protection, but fewer pups treated with the KA variant survived (**Figure 5B**). HDIT102, despite binding to an epitope overlapping with that of Hu2c, exhibited a profile more similar to HSV8: pups treated with the EG variant had significantly increased survival compared to those treated with WT, which provided similar protection as the KA variant (**Figure 5C**).

**Figure 5:**
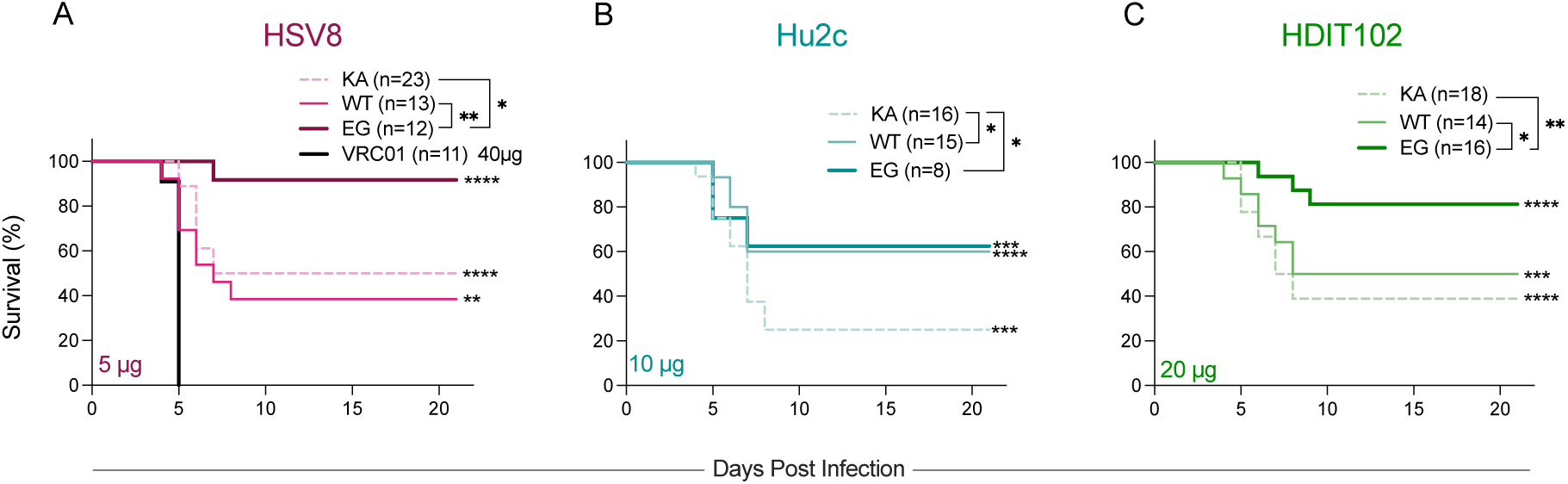
Clinical isolates are sensitive to mAb-mediated complement activation. **A-C.** HSV-specific mAbs were delivered intraperitoneally to 2-day old pups before lethal intranasal challenge with 1x10^4^ plaque forming units (PFU) of the clinical isolate HSV-1 NS. Survival of pups given gD-specific HSV8 (**A**), or gB-specific mAbs Hu2c (**B**), or HDIT102 (**C**) at indicated dose. WT, KA, or EG variants are shown in in dark thick, medium thin, light dotted lines, respectively. Statistical significance compared to WT VRC01 isotype control (black, panel A) as determined by the log-rank (Mantel-Cox) test is shown on line graphs, and inset reports the number of animals per group and statistical significance between WT, KA, and EG mAbs (*p < 0.05, **p < 0.01, ***p < 0.001, ****p < 0.0001).

Collectively, these results indicate an important contribution of complement activity in WT mAb-mediated protection afforded by Hu2c and the potential to improve antibody efficacy by enhancing antibody-complement interactions for HSV8 and HDIT102. Thus, these results extend the evidence base supporting the value of complement-mediated activities to contribute to the efficacy of both gB- and gD-specific HSV-specific mAbs to clinical isolates. They also demonstrate that the impact of enhancing or abrogating this activity for a given mAb can be virus-dependent: whereas WT and KA forms of Hu2c provided similar protection from lethal challenge with strain 17 and EG was superior, WT and EG provided similar protection from strain NS and KA was inferior.

### Improved efficacy of engineered mAbs depends on host C1q expression

To evaluate the mechanism whereby KA and EG mutations impact antibody-mediated antiviral activity *in vivo*, C1q-deficient (*C1q^-^/^-^*) pups were treated with mAbs intraperitoneally and challenged intranasally with 1x10^4^ PFU of HSV-1 NS (**Figure 6**). Improved survival following HSV-1 challenge was seen for all three Fc variants of HSV8 (**Figure 6A**), Hu2c (**Figure 6B**), and HDIT102 (**Figure 6C**) compared to a higher dose of the isotype control mAb VRC01. However, in the context of a C1q knockout host, the impact of either KA or EG mutations was lost. For gD-specific HSV8, HSV-infected mice treated with KA or EG variants survived similarly well as those treated with WT (**Figure 6A**). Likewise, for both gB-specific Hu2c and HDIT102, KA and EG variants provided equivalent protection to infected pups (**Figure 6B-C**). Altogether, these results demonstrate that the increased protection provided by the EG variants, and, perhaps perplexingly, the decreased protection provided by KA variants, was dependent on C1q expression in the host, and thus specifically mediated via the classical pathway, regardless of antigen target.

**Figure 6:**
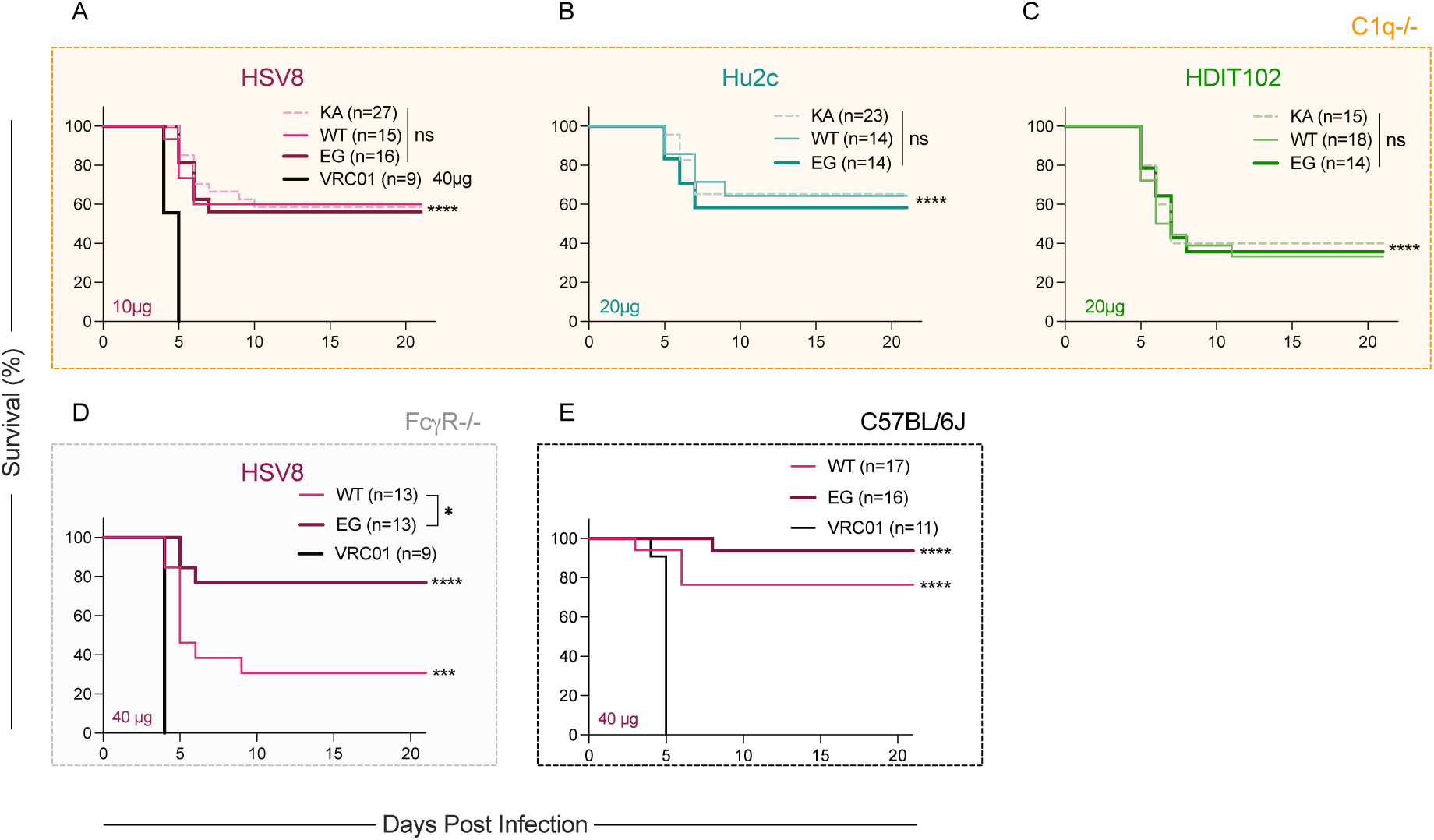
Enhanced protection afforded by the EG variant is lost in C1q-/- but maintained in FcγR-/- mice. **A-C.** HSV-specific mAbs were delivered intraperitoneally to 2-day old C1q-/- pups before lethal intranasal challenge with 1x10^4^ plaque forming units (PFU) of the clinical isolate HSV-1 NS. Survival of pups given gD-specific HSV8 (**A**), or gB-specific mAbs Hu2c (**B**), or HDIT102 (**C**) at indicated dose. **D-E**. Survival of 2-day old FcγR-/- (**D**) or C57BL/6J (**E**) pups treated with 40 μg HSV8 mAb intraperitoneally before lethal intranasal challenge with 1x10^4^ plaque forming units (PFU) of HSV-1 NS. WT, KA, or EG variants are shown in in dark thick, medium thin, light dotted lines, respectively. Statistical significance compared to WT VRC01 isotype control (black, panel **A**) as determined by the log-rank (Mantel-Cox) test is shown on line graphs, and inset reports the number of animals per group and statistical significance between WT, KA, and EG mAbs (*p < 0.05, **p < 0.01, ***p < 0.001, ****p < 0.0001).

In contrast, the enhanced efficacy of HSV8 EG relative to WT was maintained in FcγR-/- pups (**Figure 6D**) and was reduced relative to protection in the C57BL/6J background in which both knockout strains were generated (**Figure 6E**). Collectively, evaluation of host genetic factors implicates the classical complement pathway, as initiated by C1q, in the improved protection afforded by the HSV8 EG variant, which can help compensate for the loss of other antibody effector functions, such as those afforded by FcγR.

### Complement contributes to protection against HSV-1 challenge in adult mice

The immune system, including the complement cascade, is known to vary during development and with aging. In mice and humans, cascade components have been reported to have developmentally distinct levels of complement activity^59–65^. To evaluate whether antibody-dependent complement-driven mechanisms of antiviral activity contribute to HSV-1 protection in adulthood, we challenged adult (6-7 weeks of age) C57BL/6J mice intranasally with 5x10^6^ PFU of HSV-1 McKrae^66^ following intraperitoneal mAb administration (**Figure 7A**). Both HSV8 WT and KA provided protection against HSV-1 compared to the control mAb VRC01. Adult mice treated with the KA variant, however, had substantially reduced probability of survival compared to WT, demonstrating an important role for complement in mAb-mediated protection across the lifespan.

**Figure 7:**
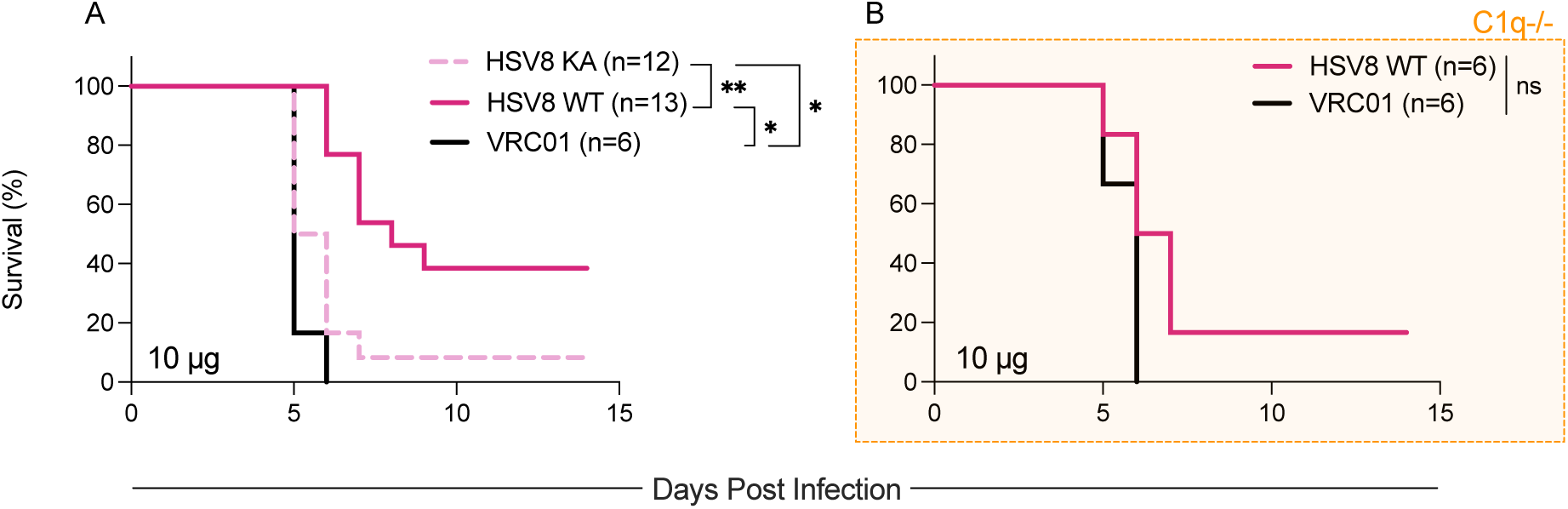
Complement abrogation results in decreased protection against HSV-1 challenge in adult mice. **A-B.** Adult (6-7 week old) C57BL/6J (**A**) or C1q-/- mice (**B**) were given 10 μg mAb intraperitoneally immediately before lethal intranasal challenge with 5x10^6^ PFU of HSV-1 McKrae. Inset reports the number of animals per group and statistical significance as determined by the log-rank (Mantel-Cox) test between indicated treatment groups (*p < 0.05, **p < 0.01).

As an orthogonal test, C1q-/- adult mice were treated with HSV8 WT and challenged intranasally with 5x10^6^ PFU of HSV-1 McKrae (**Figure 7B**). In C1q knockout mice, the same dose of WT HSV8 did not improve survival as compared to the VRC01 control mAb, suggesting that protection of adults is also dependent on the classical complement pathway.

## Discussion

Despite effort to harness the therapeutic potential of antibodies against HSV, there is yet to be approval of an effective mAb-based treatment. The value of Ab-mediated protection in both preclinical adult and neonate models of HSV infection has been seen via reduction of viral titers and increased survival^24,28,45,67–70^. However, to date, the mAbs to be clinically translated have been selected based primarily on neutralization potency and breadth. More recent work has demonstrated that these mAbs nonetheless depend strongly on effector functions to achieve optimal efficacy in preclinical models. Yet, while antibodies can be neutralizing and/or FcγR-activating, they also have the potential to activate the classical complement cascade via formation of an antigen-antibody complex and subsequent binding of C1q. Built on exquisite structural biology studies^71,72^, and with recent clinical translation of antibodies with complement-modifying mutations^46,73^, appreciation of the complement system’s role in antibody-based prevention and treatment of disease has grown. Thus, investigating the contribution that complement plays in antibody-mediated protection from HSV-1 infection, including its potential to enhance antibody-dependent antiviral activity, has prospects to lead to new preventative and therapeutic strategies.

In this work, we sought to determine the role of complement-mediated functions in antibody-dependent antiviral activity against HSV-1 infection. A panel of gD- and gB-specific antibodies was engineered to exhibit either complement-enhancing or complement-abrogating phenotypes. No differences were seen in binding to recombinant antigen between WT and Fc variants, while expected differences in C1q binding and C3b deposition were observed. Generally, the KA variants eliminated C1q binding and complement deposition while the EG variants presented an increase in these activities compared to WT. However, the degree of differences, particularly between WT and EG variants were dependent on antigen density. At a lower density, differences were sometimes more easily observed between WT and EG variants. This observation suggests that at a higher antigen density, targets are closer together, thus allowing WT IgG1 to oligomerize as efficiently as EG variants. Similar results have been observed for complement-engineering HIV-specific antibodies and antigen density in *in vitro* assays^74^.

Further, complement-dependent enhancement of viral neutralization has been reported for several viruses, including herpesviruses^52,53,75,76^. While we did see an increase in neutralization for pooled polyclonal seropositive human sera in the presence of complement, no observable difference was seen for most of the HSV-specific mAbs tested. Only CH42 (gD-specific) and BMPC23 (gB-specific) displayed complement-dependent neutralization at a high level (25%) of complement active serum. The complement-dependent increase in neutralization for these two otherwise non-neutralizing mAbs may be due to virolysis, as all components for MAC formation are present in the assay. However, we did not use C9-depleted serum to determine if lysis was the basis for increased neutralization, or if initiation of the cascade improves neutralization through steric hindrance or other means of interference with cellular attachment or entry.

Overall, *in vitro* neutralization results were not indicative of *in vivo* protection. For the gD-specific antibodies, there were varying levels of protection afforded by modulated antibody-complement interactions in *in vivo* studies. For one antibody (HSV8), there was increased survival in pups treated with the EG variant. For another (E317), there was no increased survival afforded by enhancing the interaction with C1q, but reducing it led to decreased survival of pups. Interestingly, for the non-neutralizing antibody (CH42), there was no difference in protection between WT and Fc variants even though *in vitro* data for the EG variants seemed promising (increased C1q binding, increased C3b deposition, and increased neutralization in the presence of complement). It could be that differences in epitope affect the accessibility of C1q and subsequent complement cascade proteins building upon the immune complex, or that activities against infected cells, which were not evaluated here, were more important that those directed against virions for this mAb.

Although the gB-specific mAbs included in this panel target different domains of the fusion protein, all appeared to benefit from enhanced antibody-complement interactions. The EG variants of Hu2c, HDIT102, D48 and BMPC23 resulted in increased survival of pups (compared to WT) when challenged with a lab strain of HSV-1 (strain 17). Similar superiority was generally seen in challenges with a clinical isolate, but Hu2c displayed discrepant results between clinical and lab strain, which may be attributable to differences in pathogenesis or neurovirulence between the two strains, among other factors.

In sum, differences in protection were seen among epitopes within a glycoprotein and between different glycoproteins. While differences in the rank order of protection were generally consistent, the degree of enhancement of protection afforded by EG mutation or loss triggered by KA substitution varied in association with challenge virus. Compared to the gD-specific mAbs, all the gB-specific mAbs were more conducive/sensitive to enhanced complement-mediated antiviral activity. While only two HSV glycoproteins were studied in this work, the differences seen between gD and gB suggest other surface glycoproteins may also have variable susceptibility to modulated complement activity.

Limitations of this work include that *in vitro* work was performed using a single clinical isolate of HSV-1, modeling entry into simian epithelial kidney cells. We utilized a specific set of mAbs at set concentrations and in the presence of a limited dilution series of guinea pig serum, which often exhibits higher activity than human or murine serum, as the source of complement. We did not measure the relative contributions of effector functions known to contribute to *in vivo* efficacy for many of these mAbs^24,45,70^. While EG and KA mutations are reported to exhibit no effect on FcγR binding, activation of the cascade can lead to the production of opsonins which tag immune complexes for phagocytic cells^10^. Anaphylatoxins (C5a and C3a) resulting from the cascade can also contribute to the mechanism of action by attracting effector cells with complement receptors to the area of infection^5,77^. These activities may have contributed to the increased *in vivo* protection seen with the EG variant of some mAbs. Our *in vitro* work did not suggest that neutralization was the main mechanism through which complement contributed to antibody-mediated antiviral activity. However, plaque assays were performed using diluted guinea pig serum. Complement-mediated neutralization could play a larger role *in vivo* where complement regulators are present and soluble complement proteins are produced continuously. Alternatively, modification of lysis of virions and/or infected cells via MAC formation may be the sole contributing mechanism of action of the KA and EG variants. Given the sensitivity of *in vitro* complement assays to test variables^78,79^, we did not try and thus cannot further pinpoint the mechanism(s) of action by which the EG variants provided increased protection.

*In vivo* work was performed in one mouse strain with a single laboratory strain and a single clinical isolate of HSV-1, with one route of challenge, mode of mAb administration, timed simultaneously. While complement proteins are highly conserved between humans and mice, and their functional roles are similar despite some sequence and expression level variation ^80–82^, we presume that our design choices are likely to influence both magnitude and mechanism of protection. Experimental variation in mechanism of action against different viruses and sites and levels of viral inoculum exposure is consistent with evolutionary pressure to provide broad protection across genetically diverse hosts. Systematic collaboration for host defense across differing anti-viral activities has been proposed explicitly for both HIV- and SARS-specific antibodies^83^. Consistent with this hypothesis, while better *in vitro* function was generally associated with better *in vivo* efficacy, direct and linearly quantitative relationships between *in vitro* potency of any single activity assessed and *in vivo* efficacy were not observed in our study.

We focused on HSV-1 as it is the most prevalent subtype globally and in the United States^84^. Further work needs to be done to determine if findings translate to HSV-2 given that gC differs both structurally and functionally between HSV serotypes^85^. While the virus’ anti-complement adaptations may differ, we expect that an antibody therapeutic manipulating complement would serve a majority of the population, as there is only a 0.03%^86,87^ estimated prevalence of genetic complement deficiencies. Though studied in a neonatal model here, epidemiologic evidence^88–91^ suggests than enhancing antibodies beyond their natural abilities is not likely to necessary for a mAb to provide robust protection of neonates, who benefitted from enhancement despite reports of lower levels of cascade components than adult mice^92^, a phenotype also observed in humans^93,94^. Instead, functional enhancement may be more likely to contribute clinically to mAbs for reducing chronic reactivation, which is observed in some individuals despite robust seroconversion and neutralizing activity^95–97^. The recently reported failure^98^ of Hu2c (HDIT101) to reduce the percentage of days participants experienced lesions, but the marginally extended time to recurrence and statistically significant reduction in reactivation rate, present hope that mAb optimization, such as offered by Fc engineering, may lead to new clinical options.

Overall, our data demonstrates the importance of complement for some mAbs. Yet, the ability of each mAb to mediate protection from lethal challenge was variably impacted by Fc domain modifications that eliminated or enhanced Fc domain interactions with the cascade initiator C1q. Abrogating complement activity reduced protection afforded by some mAbs, but enhancing C1q deposition improved efficacy for a greater number of mAbs, including all four gB-specific mAbs tested. These profiles were also dependent on the viral strain and host genetics. In sum, these findings demonstrate that optimal protection against HSV likely requires diverse antibody functions, including complement-mediated activities, and provide insight into the clinical prospects of antibody-based therapeutics by highlighting functional differences among antibodies and target antigens and across host and pathogen.

## Methods

### Sex as a biological variable

Male and female mice were used in HSV-1 challenge experiments. Sex was not considered as a biological variable.

### Cells

Vero cells (ATCC: CCL-81) were maintained in DMEM containing 10% FBS and 1% penicillin/streptomycin at 37C with 5% CO2. EXPI293F cells (ThermoFisher) were maintained in Expi293F Media (Thermo Fisher) in a reach-in CO2 incubator at 37°C with 8% CO2 on an Innova 2300 platform shaker at 125 RPM.

### Animals

Male and female C57BL/6J (RRID: IMSR_JAX:000664) and B6 (Cg)- *C1qa^tm1d(EUCOMM)Wtsi^*/TennJ (C1q−/−) (RRID:IMSR_JAX:031675) mice were purchased from The Jackson Laboratory or bred in animal facilities at Dartmouth College in accordance with institutional animal care and use committee (IACUC) protocols (Dartmouth College IACUC 2151). Male and female B6.129P2-Fcer1gtm1Rav N12 (FcyR−/−) (model: 583) were purchased from Taconic Labs. Offspring of both sexes were evaluated in challenge studies.

### Mouse procedures and viral challenges

Antibodies were administered to 2-day old pups under 1% isoflurane anesthesia via the intraperitoneal route with a 25 μL Hamilton syringe in a volume of 20 μL. Pups were immediately infected intranasally with 1x10^4^ PFU of HSV-1 strain 17 (Brown et al.)^56^ or 1x10^4^ PFU of HSV-1 strain NS (provided by H. Friedman)^58^ in a volume of 5 μL. Viral stocks were prepared using Vero cells as previously described^99,100^. Over a 21 day follow up period, pups were monitored and weighed daily. Endpoint criteria for survival studies was defined as excessive morbidity (hunching, spasms, and/or paralysis) and/or >10% weight loss from the previous measurement. For adult mice, 6-7 week old mice of both sexes were administered antibodies via intraperitoneal route with a 1mL Leur lock insulin syringe in a volume of 200 μL. Mice were immediately infected intranasally with 5x10^6^ PFU of HSV-1 McKrae under 1% isoflurane anesthesia. Over a 15 day follow up period, mice were monitored and weighed daily. Endpoint criteria for adult mice was defined as excessive morbidity (hunching, spasms, and/or paralysis) and/or weight loss >15%.

### Monoclonal antibodies

E317, HSV8, CH42, Hu2c, HDIT102, D48, and BMPC-23 VH and VL sequences were collected from published sequences or published structures^28,37,45,101^ and synthesized by Twist Biosciences as full length IgG1 heavy chains and Kappa (E317, HSV8, CH42, Hu2c,BMPC-23) or Lambda (HDIT102) constant light chains. Recombinant DNA was cloned into pCMV expression vectors (Addgene). Complement-enhancing and abrogating mutations (E430G and K322A, respectively)^33,34^ were introduced into the IgG1 heavy chain backbones via QuickChange Lightning Site-Directed Mutagenesis (Agilent) according to the manufacturer’s instructions using mutagenic primers (KA_forward: GCAAGGAGTACAAGTGCGCGGTCTCCAACAAAGCCC; KA_reverse: GGGCTTTGTTGGAGACCGCGCACTTGTACTCCTTGC; EG_forward: CTCCGTGATGCATGGGGCTCTGCACAACC; EG_reverse: GGTTGTGCAGAGCCCCATGCATCACGGAG). Heavy and light chain expression plasmids were sequence confirmed via whole plasmid sequencing (Azenta/GENEWIZ). Antibodies were expressed through co-transfection of heavy and light chain plasmids in Expi293 HEK cells (Thermo Fisher) according to the manufacturer’s instructions. Six days after transfection, cultures were spun at 3750 rpm for 1 hr to pellet the cells, and supernatants were filtered (0.45 μm). IgG was affinity purified using a custom-packed 2 mL protein A column (Cytiva) and eluted with 100 mM glycine pH 3, which was immediately neutralized with 1 M Tris buffer pH 8. The eluate was then concentrated and buffer exchanged into PBS using Amicon 30kDa centrifugal filters. mAbs were passed over endotoxin removal columns (ThermoFisher) according to the manufacturer’s instructions and aliquoted before storage at -80°C.

### Isolation of polyclonal IgG

Human serum samples were analyzed for HSV seropositivity using a bead-based assay to detect for the presence of gD/gB specific antibodies. Three individual donor samples with high titers were pooled together, and IgG was isolated using a Melon Gel IgG purification kit (ThermoFisher). The flowthrough was concentrated and buffer exchanged into PBS using Amicon 30kDa centrifugal filters.

### Viral neutralization

Viral neutralization of HSV-1 was performed via plaque reduction neutralization as previously described^70,102^. Briefly, serial dilutions of mAb or polyclonal IgG were incubated with 100 PFU of HSV-1 st17 at 37°C for 1 hr before being added to confluent Vero cell monolayers grown in 6 well plates (Corning) and incubated for 1 hr at 37°C with shaking every 15 minutes to allow infection. Afterwards, a methylcellulose (0.02g/mL) overlay was added to the cells and plates were incubated for 72 hr at 37°C with 5% CO2, at which time the overlay was removed, and cells were fixed with 1:1 methanol:ethanol for 30 min at RT. Plates were then stained with 12% Giemsa overnight. The stain was removed, and plaques were manually counted on a light box. Viral neutralization (%) was calculated as [(number of plaques in virus only control well - number of plaques counted at mAb dilution)/# of plaques in virus only control well]*100. Assays were performed in technical replicates. Cells only and virus only (no mAbs) wells served as controls. To measure antibody-dependent complement-assisted neutralization, a 1:4 or 1:64 dilution of guinea pig serum (Cedarlane) was added the virus:mAb incubation. A complement only (virus present, no mAbs) well served as a control for antibody-independent viral inhibition, along with heat-inactivated complement serum (56°C, 30 min) well as a negative control for baseline levels. Assays were performed in technical and biological duplicate.

### Antigen binding

Antigen beads were prepared by coupling HSV-1 gD or gB to coded Magplex superparamagnetic carboxylated magnetic microparticles (Luminex Corp) using carbodiimide cross-linking chemistry as previously described^103^. Antigen beads were diluted in assay buffer to 500 beads/well and then incubated with serially diluted antibody for 2 hrs at RT with shaking. Following antibody binding, the plate was washed with assay buffer (PBS with 0.01% BSA and 0.005% Tween 20) using an automatic plate washer and bound antibody was detected with a PE-conjugated mouse anti-human IgG Fc (Southern Biotech) for 1hr at RT with shaking. The plate was then washed again and beads were resuspended in 50 µL of sheath fluid (Luminex Corp). Data was acquired on a Luminex FLEXMAP 3D Instrument System which detects beads and measures PE fluorescence to calculate the MFI level of each analyte. Assays were performed in technical and biological duplicate.

### Measurement of binding of C1q to mAbs

Antibody-opsonized antigen beads were prepared as described above and incubated with 0.1 µg/well (1.0 µg/mL) biotin anti-human C1q (ThermoFisher) for 1 hr at RT with shaking. After 1 hour, the plate was washed using an automatic plate washer and incubated with Strep-PE reagent (1ug/ml; Agilent) for 30 minutes. The plates was then washed using an automatic plate washer and beads were resuspended in 50 µl of sheath fluid. Data was acquired on a Luminex FLEXMAP 3D Instrument System which detects beads and measures PE fluorescence to calculate the MFI level of each analyte. Wells with no mAb were used as a control. An HIV-specific IgG1 (VRC01) was used as a negative control. Assays were performed in technical and biological duplicate.

### Antibody-dependent complement deposition

Antibody-dependent Complement Deposition (ADCD) was evaluated by treating antibody-opsonized antigen beads prepared as described above with 1:100 dilution of guinea pig complement serum (Cedarlane). The plate was sonicated and vortexed to allow for mixing of beads and antibodies before being sealed and incubated at 37°C for 30 min and washed with assay buffer. For C3b detection, 40 µL of 0.65 µg/mL of biotin-conjugated anti-guinea pig C3 (ICL, GC3-60B-Z) was added to the plate and incubated for 40 minutes at RT with shaking. This was followed by an additional 20ul of 1ug/mL Strep-PE (Agilent) and incubated for 20 minutes at RT with shaking before a final wash, resuspension, and data acquisition as described above. Heat-inactivated guinea pig serum (heated for 30 minutes at 56°C) was used as a negative control. An HIV-specific IgG1 (VRC01) was used as a negative control. Complement only (no mAbs) and heat-inactivated complement served as controls for antibody-independent activity and baseline levels. Assays were performed in technical and biological duplicate.

### Structural analysis

The crystal structure of gD (PDB: 1JMA^104^) was colored to indicate mAb epitopes as defined by prior studies (E317^36^ PBD: 3W9E; CH42 from Wang el al.,^28^; HSV8 from De Logu et al.,^37^). The crystal structure of the pre- (left, PDB: 6z9m^105^) and post- (right, PDB: 2gum^106^) fusion structures of the gB trimer were colored to indicate mAb epitopes as defined by prior studies (Hu2c^31^ PBD: 8RGZ; HDIT102^31^ PDB: 8RH0; D48^107^ PDB: 8KFA; BMPC-23^23^ PDB: 7UHZ).

### Statistics

Structural visualization was performed using ChimeraX. Data visualization and statistical analysis was performed using Prism version 10.6.1 (GraphPad Software). Log rank tests were used to assess statistical significance in survival experiments.

### Study approval

Animal procedures were performed in accordance with Dartmouth’s Center for Comparative Medicine and Research policies and following approval by Dartmouth’s Institutional Animal Care and Use Control (Protocol number: 00002151).

## Data availability

Study data are available in the **Supporting Data Values File**.

## Acknowledgements

We thank all members of the Ackerman and Leib laboratories for invaluable feedback and scientific discussion. Molecular graphics and analyses were performed with UCSF ChimeraX, developed by the Resource for Biocomputing, Visualization, and Informatics (RBVI) at the University of California, San Francisco, with support from National Institutes of Health R01-GM129325 and the NIAID Office of Cyber Infrastructure and Computational Biology. Figure 4 contains an image created in BioRender. Ackerman, M. (2027) https://BioRender.com/8lb4ige

