## Supplemental for "Interaction with complement factor C1q can contribute to and improve monoclonal antibody-mediated protection against lethal HSV-1 challenge"

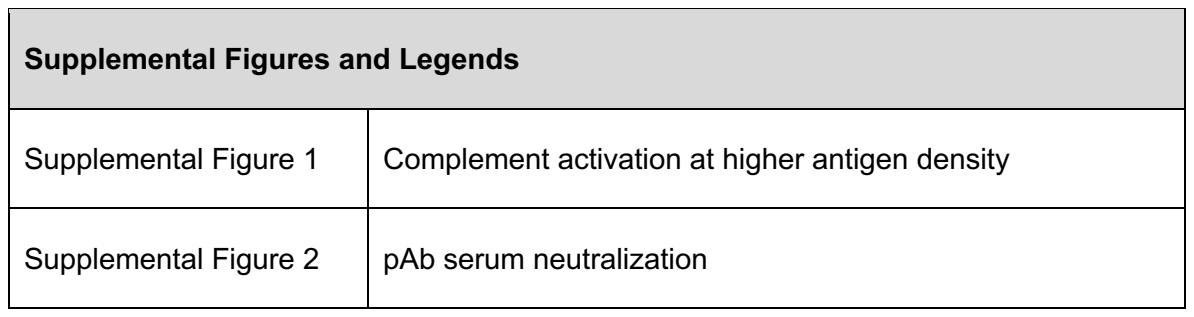

| Supplemental Figures and Legends |  |
| --- | --- |
| Supplemental Figure 1 | Complement activation at higher antigen density |
| Supplemental Figure 2 | pAb serum neutralization |

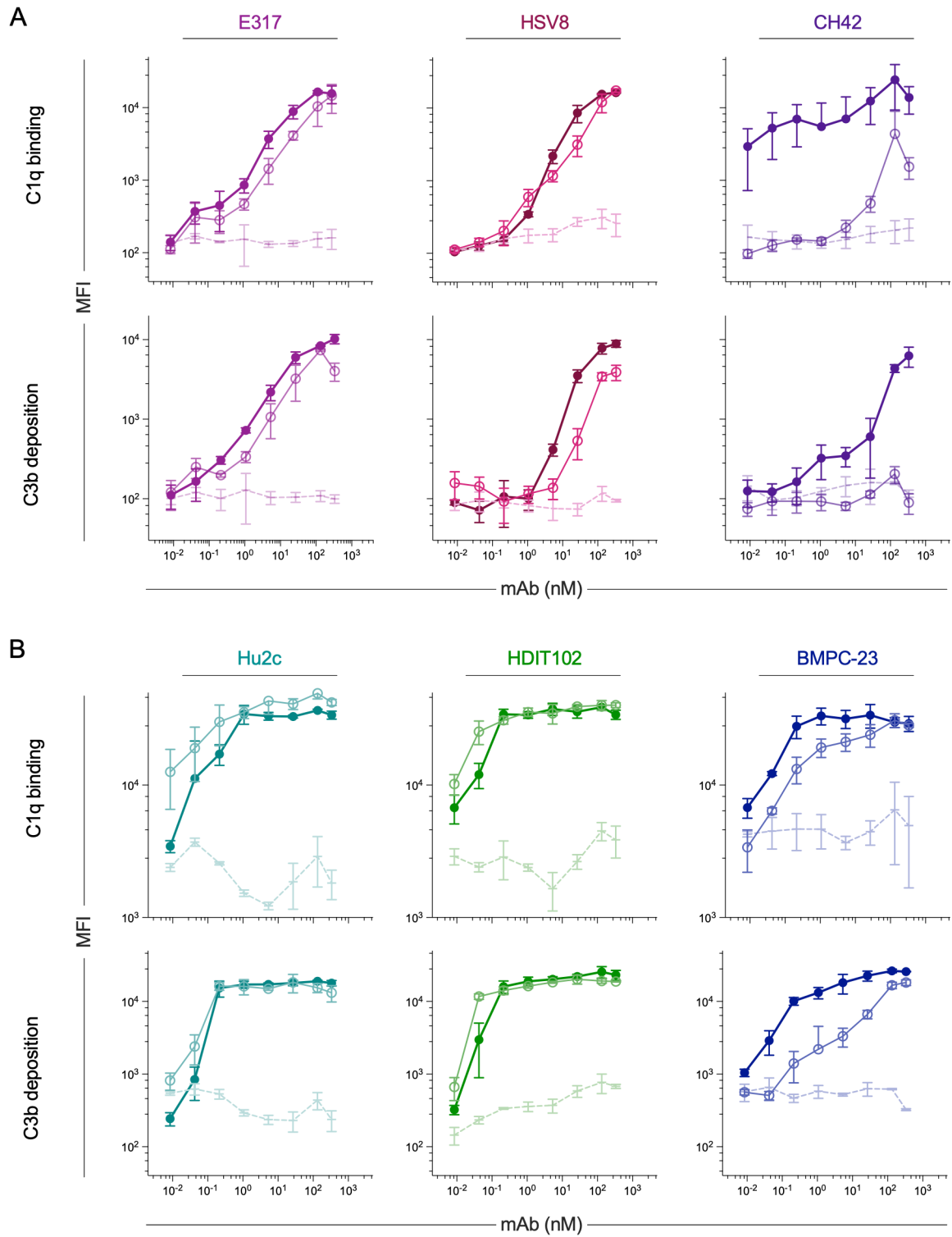

**Supplemental Figure 1: Complement activation at higher antigen density. A-B.** Functional characterization of indicated gD- (**A**) and gB (**B**) specific mAbs. C1q binding (top rows) and C3b deposition (bottom rows). Graphs depict median fluorescent intensity (MFI). Error bars indicate standard deviation across replicates.

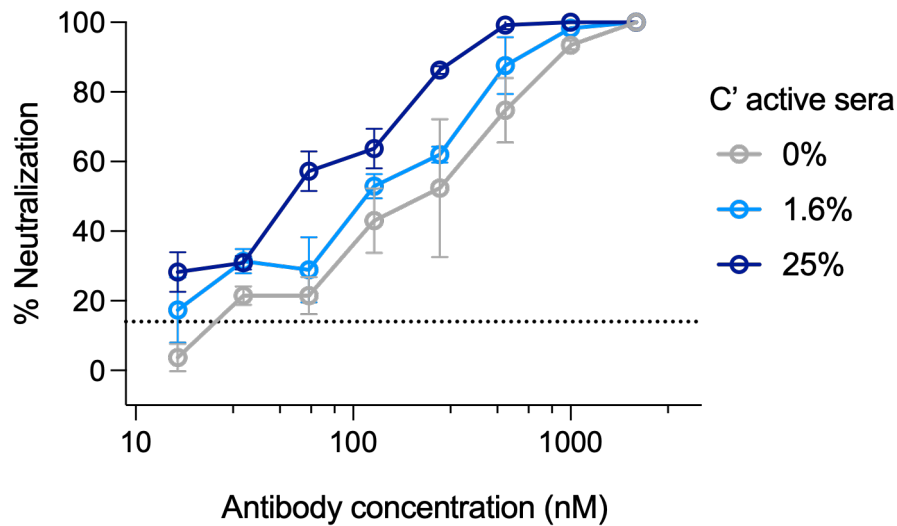

**Supplemental Figure 2: Neutralizing activity of seropositive human serum increases in the presence of complement.** Polyclonal IgGs (pAbs) were isolated from pooled seropositive human serum. pAbs were titrated and assessed for neutralization the absence (gray) and presence of complement at a high (dark blue) and low (light blue) dilution. Active complement serum (without pAbs; dotted line) was included as a negative control for baseline complement-mediated activity. Error bars indicate standard deviation.
